# A stimulus artefact undermines the evidence for independent ON and OFF channels in stereopsis

**DOI:** 10.1101/295618

**Authors:** Jenny C. A. Read, Bruce G. Cumming

## Abstract

Early vision proceeds through distinct ON and OFF channels, which encode luminance increments and decrements respectively. It has been argued that these channels also contribute separately to stereoscopic vision. This is based on the fact that observers perform better on a noisy disparity discrimination task when the stimulus is a random-dot pattern consisting of equal numbers of black and white dots (a “mixed-polarity stimulus”, argued to activate both ON and OFF stereo channels), than when it consists of all-white or all-black dots (“same-polarity”, argued to activate only one). However, it is not clear how this theory can be reconciled with our current understanding of disparity encoding. Recently, a binocular convolutional neural network was able to replicate the mixed-polarity advantage shown by human observers, even though it was based on linear filters and contained no mechanisms which would respond separately to black or white dots. Here, we show that the stimuli used in all these experiments contain a subtle artefact. The interocular correlation between left and right images is actually lower for the same-polarity stimuli than for mixed-polarity stimuli with the same amount of disparity noise applied to the dots. Since our current theories suggest stereopsis is based on a correlation-like computation in primary visual cortex, it is then unsurprising that performance was better for the mixed-polarity stimuli. We conclude that there is currently no evidence supporting separate ON and OFF channels in stereopsis.

## Introduction

In the journal *Nature* twenty-three years ago, Harris and Parker (1995) made a striking claim about stereoscopic vision. They argued for the existence of independent neural mechanisms for bright and dark information in stereopsis. Their evidence came from the performance of their two observers on a depth discrimination task made challenging by the addition of disparity noise. The stimulus was a random-dot stereogram with a vertical step edge. The mean disparity was opposite on either side of the edge, but each dot was given a noise disparity drawn from a Gaussian distribution with a given standard deviation. The task was to detect which half of the stereogram was closer. Observers performed better when the stimulus was made up of equal number of black and white dots on a gray background (Figure 1A) than when all dots were black (Figure 1B) or white (Figure 1C).

**Figure 1.**
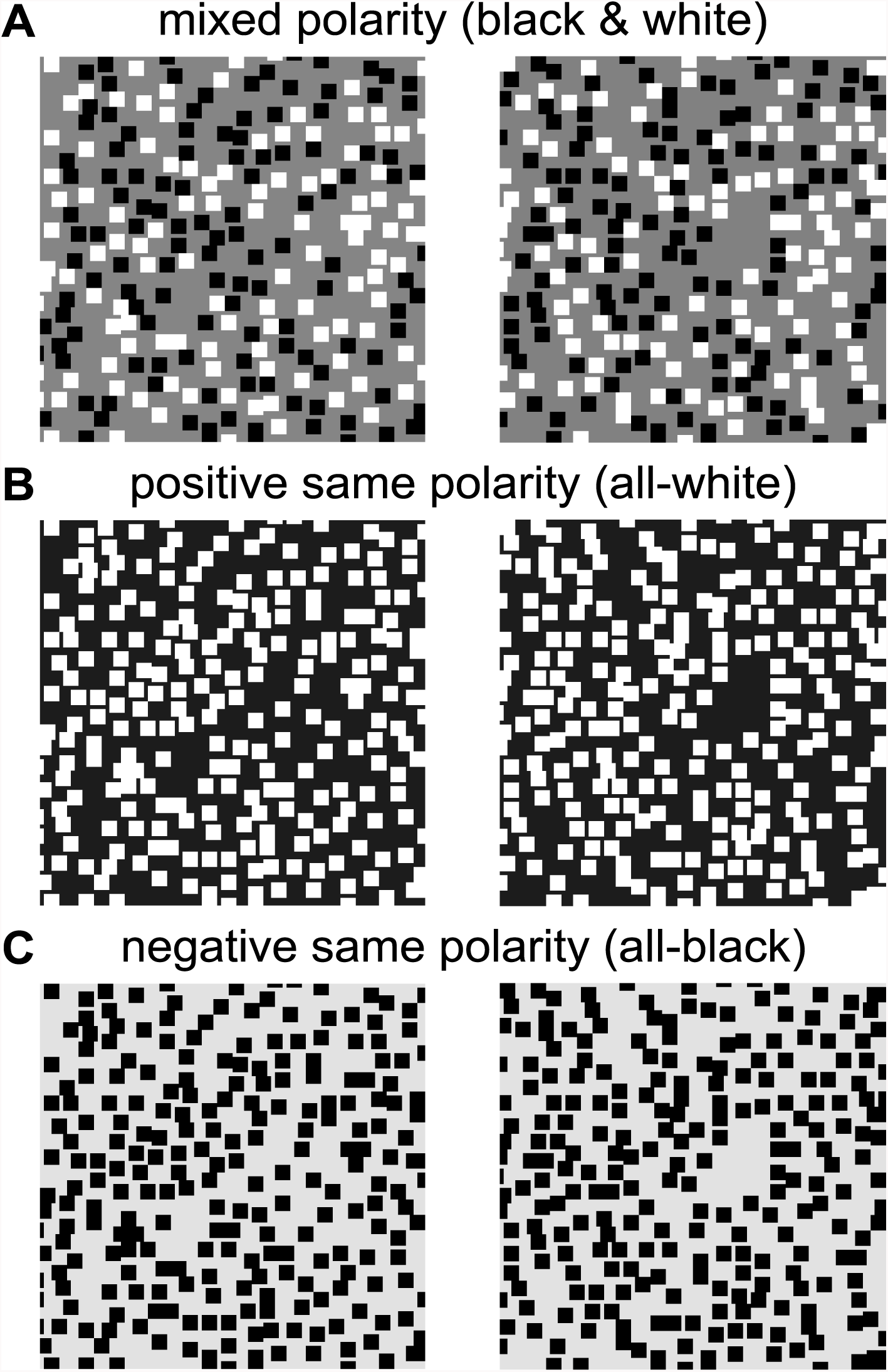
Mixed and same-polarity stimuli. The stimuli are stereograms suitable for free fusion. A: Mixed-polarity dot stimuli (black and white dots on a gray background); B,C: same-polarity stimuli with either black-only or white-only dots. All three examples are for the no-overlap condition, where dots are not allowed to overlap/occlude one another, as in Harris & Parker (1995). The images are 100×100 pixels and the dots are 4×4 pixels with density *ρ*=0.4, corresponding to an average of 250 dots per image). The images have zero mean disparity and Gaussian disparity noise with standard deviation equal to the dot size. The images have been normalized such that they all have the same mean and standard deviation of luminance. Psychophysically, the mixed-polarity advantage is unaffected by this normalization, showing that it cannot be explained by a contrast artefact.

One can imagine an ideal observer solving this task by computing the mean disparity of *N* dots on either side of the step-edge. Despite the noise, if the observer averaged enough dots on each side of the boundary, they would correctly judge the sign of the step. Harris & Parker worked out what *N* would have to be for this ideal observer to match the performance of their observers. They found that the implied number of dots was around twice as large for mixed-polarity stimuli as for same-polarity stimuli, representing a doubling of statistical efficiency. Harris & Parker related this to the problem of stereo correspondence. Before the disparity of a dot can be identified, it has to be successfully matched up with the corresponding dot in the other eye. When all the dots are the same color, each dot in one eye could potentially be the correct match for any dot in the other eye. But if the stereo system only matches black dots with black dots and white dots with white ones, the number of false matches is halved for mixed-polarity stimuli. This could enable more dots to be successfully matched up, increasing the number *N* of disparities which can be averaged on each side of the step-edge and so improving performance. They suggested that the independent processing of black and white dots could be mediated by the separate ON and OFF channels that are well-established early in the visual system (Jiang, Purushothaman, & Casagrande, 2015; Schiller, 2010; Schiller, 1992).

Harris & Parker’s mixed-polarity advantage was replicated with several more observers by Read, Vaz and Serrano-Pedraza (2011). These authors explored a different way of making the task hard: they made a certain proportion of the dots uncorrelated. That is, some dots had the disparity associated with the set-edge, while other dots were removed and then replaced at random, independently in each eye. Read et al argued that presented even more of a challenge to stereo correspondence. In the disparity-noise version of the task, each dot does have a definable disparity, even if this may be hard for the visual system to extract. But in the decorrelated version of Read et al, the uncorrelated dots do not have a disparity at all. Read et al found that the mixed-polarity advantage was even more pronounced in the decorrelation version of the task, with an implied efficiency ratio of up to five. Although this is not as neat as the original doubling of efficiency, mapped on to distinct ON/OFF channels, it is still consistent with Harris & Parker’s argument that stereo correspondence is easier in mixed-polarity stimuli.

The problem is that it is not at all clear how to reconcile this with the understanding of disparity encoding which has developed since Harris & Parker’s 1995 paper was published. Physiologically, stereo correspondence is believed to begin in primary visual cortex (V1). Many V1 neurons are tuned to disparity in random-dot patterns like those shown in Figure 1. A few years before Harris & Parker’s paper, Ohzawa, DeAngelis and Freeman (1990) had published a landmark paper in which they introduced a simple mathematical model describing how these cells could encode disparity. In the quarter-century since then, this binocular energy model has become the canonical description of the early stages of disparity encoding. It has been extensively tested against real V1 neurons and although modifications are certainly required, the basic principle underlying the energy model has been vindicated (Cumming & DeAngelis, 2001; Henriksen, Tanabe, & Cumming, 2016; Ohzawa, 1998; Read, 2005); notably, the model has made successful predictions about the response of real neurons to impossible stimuli (Cumming & Parker, 1997). Indeed, stereopsis has become one of the areas where we have the clearest understanding of how perceptual experience relates to early cortical encoding (Parker, 2007; Read, 2014; Roe, Parker, Born, & DeAngelis, 2007).

However, the energy model does not recognise image features such as dots. Rather, it effectively computes the interocular cross-correlation between left and right images after filtering with filters that are bandpass for orientation and spatial frequency (Allenmark & Read, 2011; Qian & Zhu, 1997). For the energy model, the dark spaces represented by the background in an all-white-dot stimulus should serve just as well as black dots. Recently, Ichiro Fujita and colleagues have presented psychophysical evidence which they argue implies a separate “matching” computation, which is sensitive to contrast polarity and will not match black dots with white (Doi & Fujita, 2014; Doi, Takano, & Fujita, 2013; Doi, Tanabe, & Fujita, 2011). In principle, such a matching computation is consistent with Harris & Parker’s theory, but its neural substrate is unknown. We have recently shown that the psychophysical data can be equally well explained by a slight modification (an additional output nonlinearity) to the energy model (Henriksen, Cumming, & Read, 2016). This model makes specific predictions regarding the effect of dot size and dot density on performance, which were tested and verified, and is also consistent with the properties of V1 neurons (Henriksen, Read, & Cumming, 2016). Thus, Doi and Fujita’s observations do not require a dot-matching process.

Thus it is not currently clear how one could build a model of V1 neurons which would provide a neuronal basis for Harris & Parker’s independent mechanisms. One might consider the modified version of the energy model proposed by Read, Parker, & Cumming (2002), where monocular inputs are half-wave rectified before binocular combination. This enables a binocular neuron to receive input only from an ON or an OFF channel. However, Read et al (2011) showed that in this model as for the original energy model, disparity tuning curves have the same amplitude for mixed-as for same-polarity stimuli. The reason, again, is that the energy model and variants sense interocular correlation, but do not look for specific image features. Similarly, there is not a simple mapping between white/black dots and ON/OFF channels. “OFF” detectors based on linear filters are activated by light decrements in the receptive field centre. But they are also activated by light increments in the surround. Consequently, a single-polarity pattern containing only bright dots does not stimulate only ON channels, since the OFF channels are stimulated by the dark regions between the bright dots. Similarly, in mixed polarity patterns the OFF channel is not blind to the white dots. So, even if ON and OFF channels are treated separately, they would not provide independent estimates of disparity. This is what makes the observation by Harris and Parker so striking, and so hard to reconcile with current views about disparity processing.

Very recently, Goncalves & Welchman (2017) put forward the first simulation that reproduces the mixed-polarity advantage observed psychophysically. Their model was a convolutional neural network, which they called a Binocular Neural Network. They explained its mixed-polarity advantage by noting that “the network depends on the activity of the simple units moderated by readout weights. Presenting mixed versus single-polarity stimuli increases the simple unit activity, in turn changing the excitatory and suppressive drives to complex units. We found that mixed stimuli produce greater excitation for the preferred output unit and increased suppression to the non-preferred unit”. This is puzzling, because Goncalves and Welchman’s Binocular Neural Network is simply a generalized version of the original energy model (Read & Cumming, 2017), with more subunits and a half-linear instead of half-squaring output nonlinearity. These additional ingredients could not enable the model to work as envisaged by Harris & Parker, matching dot to dot in a way sensitive to contrast sign rather than simply responding to the interocular correlation between left and right images. Despite this, the model in Gonclaves and Welchman clearly did reproduce the effect reported by Harris and Parker. Even more striking, their model reproduced a second subtle feature of human psychophysics. In constructing their stimuli, Harris and Parker did not permit dots to overlap one another. Read et al (2011) reproduced the results of Harris and Parker, but also showed that the mixed-polarity advantage is no longer present when dots were allowed to overlap. Goncalves and Welchmann reproduced this behaviour also.

It seems remarkable that a relatively simple model based on initial linear filtering can reproduce these psychophysical phenomena which seem to depend on specific image features (dots). Here, we try to understand what explains this behaviour. We will show that avoiding dot-overlap has a subtle effect on the binocular cross-correlation in the images, and that this is different for same-polarity and mixed-polarity stimuli. As a result, models much simpler than that of Goncalves and Welchman can also explain these phenomena. In these simple models, it is clear that there are not separate ON and OFF channels. As a result, the existing evidence does not support the conclusion that human stereopsis uses separate ON and OFF channels. Of course, our analysis does not prove that humans do not use separate channels either – further experimental work will be required to see if the explanation offered here and by Goncalves and Welchman correctly explains human stereopsis.

## Methods

### Pearson correlation coefficient

We measure the correlation between left and right-eye images with the standard formula for the sample Pearson correlation coefficient, *r*. The images consist of *n* pixels in each eye. *L*_*j*_, *R*_*j*_ is the value of the *j*^th^ pixel in the left, right eye. Then the sample Pearson correlation coefficient is

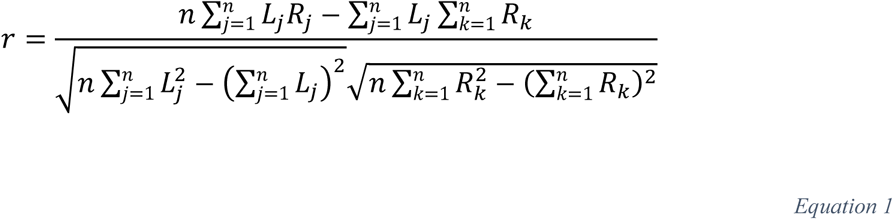

For brevity, below we will write the sums as ∑*LR* etc, omitting the indices. Equation 1 describes the correlation at zero disparity, but a similar expression holds for any uniform disparity if ∑*LR* is computed between appropriately displaced pixels. In Figure 2 and Figure 3, we plot this correlation coefficient for zero-disparity images. In Figure 4, we plot it as a function of image displacement for disparate images.

**Figure 2.**
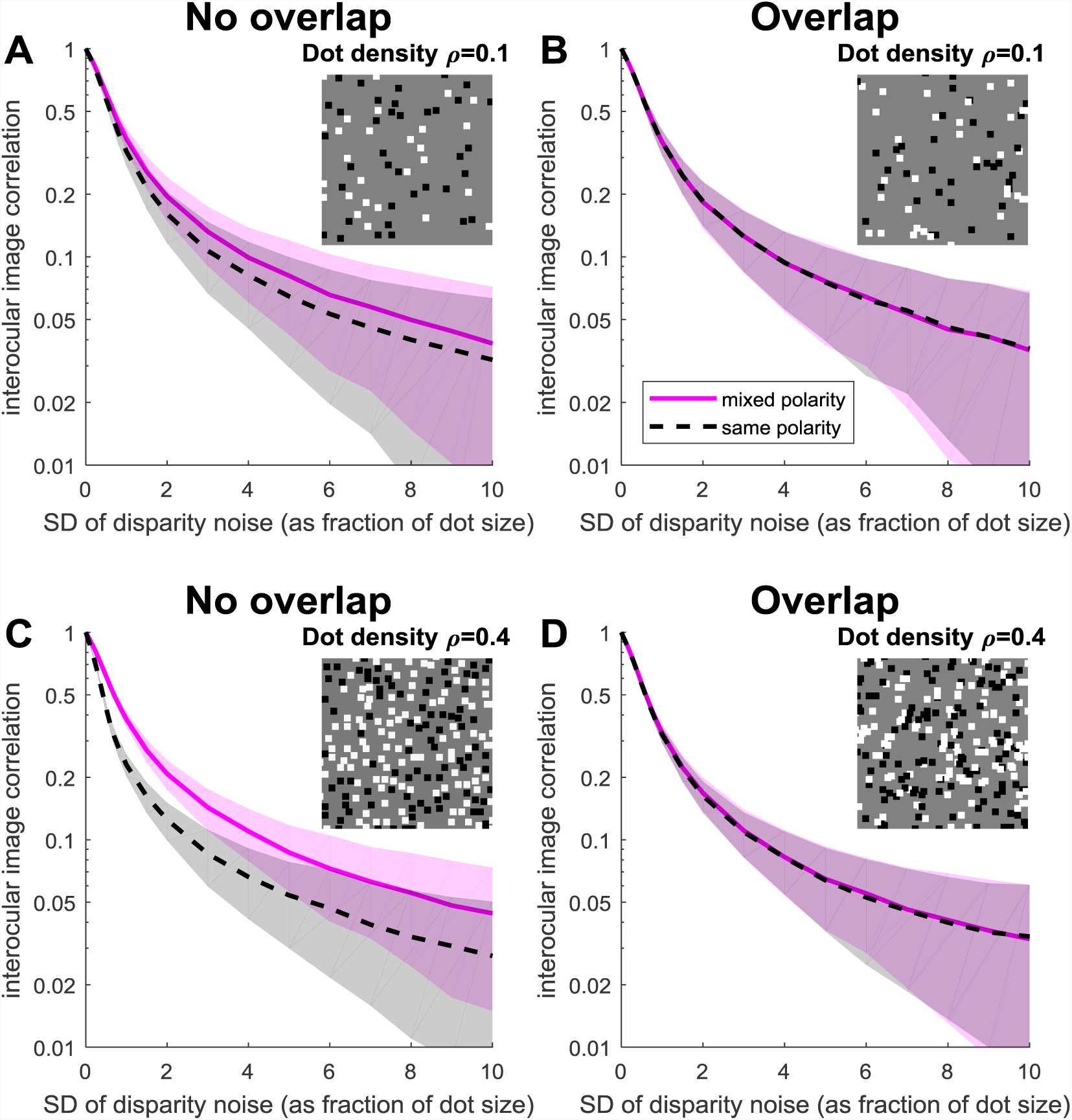
Disparity noise is more disruptive for same-polarity patterns. Interocular image correlation plotted as a function of disparity noise (the standard deviation of the Gaussian noise distribution as a fraction of the dot size in the pattern). Shaded regions show. Lines show the mean Pearson correlation coefficient between left and right images, averaged over 1000 different random-dot patterns; shaded regions show ±SD. Note that the vertical axis is logarithmic and all images have zero mean disparity. Insets show example mixed-polarity images. Same-polarity images are the same except all dots are black or all white. In the simulations, the mixed-polarity images were generated first and then the dot colors were manipulated to produce same-polarity images. Thus, the same dot patterns were used for both conditions. AC: No-overlap condition (dots are placed only in empty regions of the stimulus); BD: Overlap condition (dots are placed at random, occluding one another when they overlap). AB: Low-density (*ρ*=0.1); CD: High-density (*ρ*=0.4). The same number of dots were drawn in each case, so the area unoccupied by dots (the background) is a little higher for the no-overlap condition. Insets show example mixed-polarity images.

**Figure 3.**
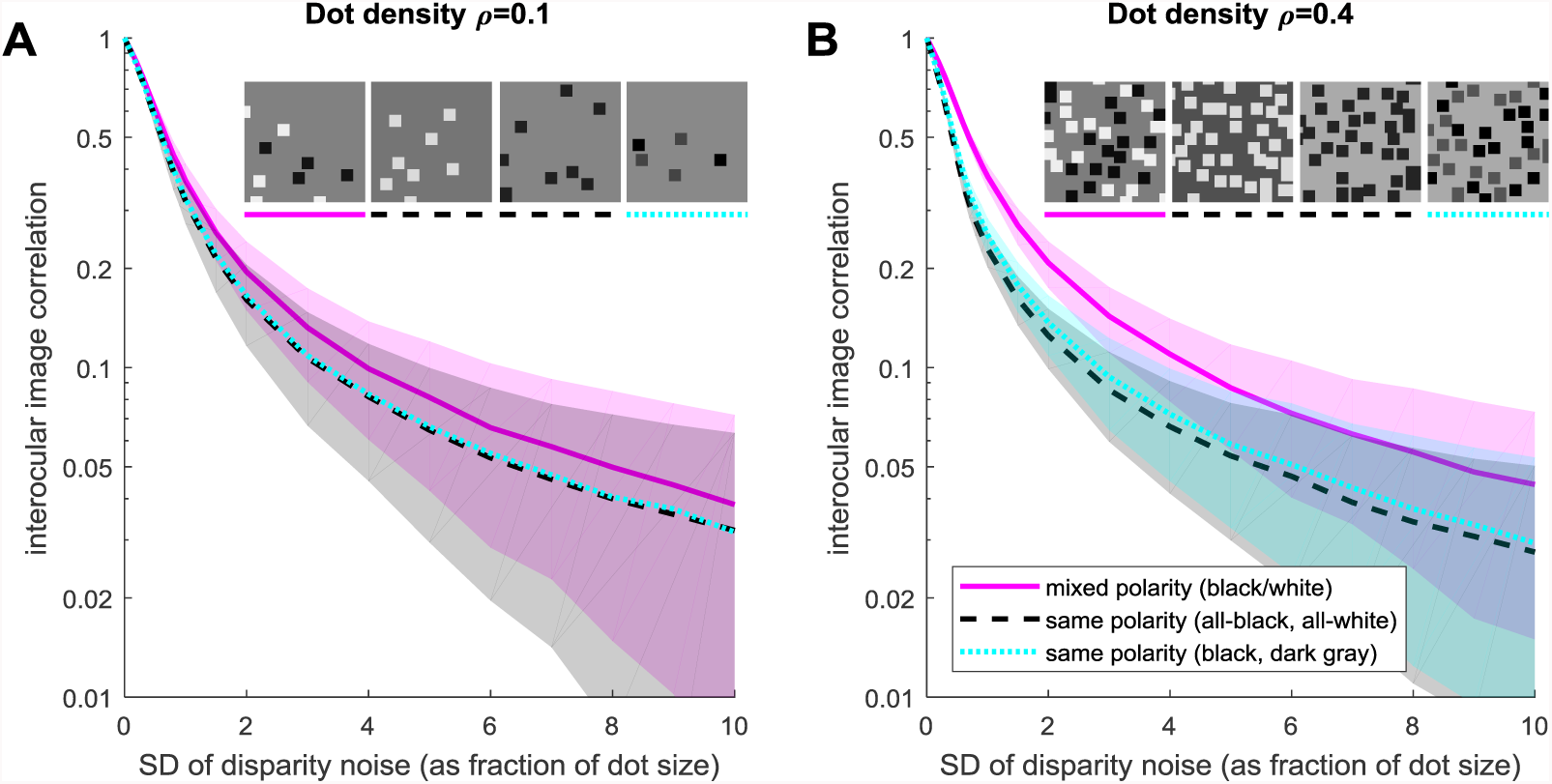
For patterns with no dot overlap, correlation is lower in same-polarity dot patterns even where the dots have two different contrasts. As Figure 2, but the cyan/dotted curves show results for same-polarity patterns with equal numbers of black and dark gray dots, for comparison with same-polarity patterns with all-black or all-white dots (black/dashed curves). There is no significant difference between the 3 same-polarity patterns. A: Low-density (*ρ*=0.1); B: High-density (*ρ*=0.4); no dot overlap throughout. Other details as in Figure 2.

**Figure 4.**
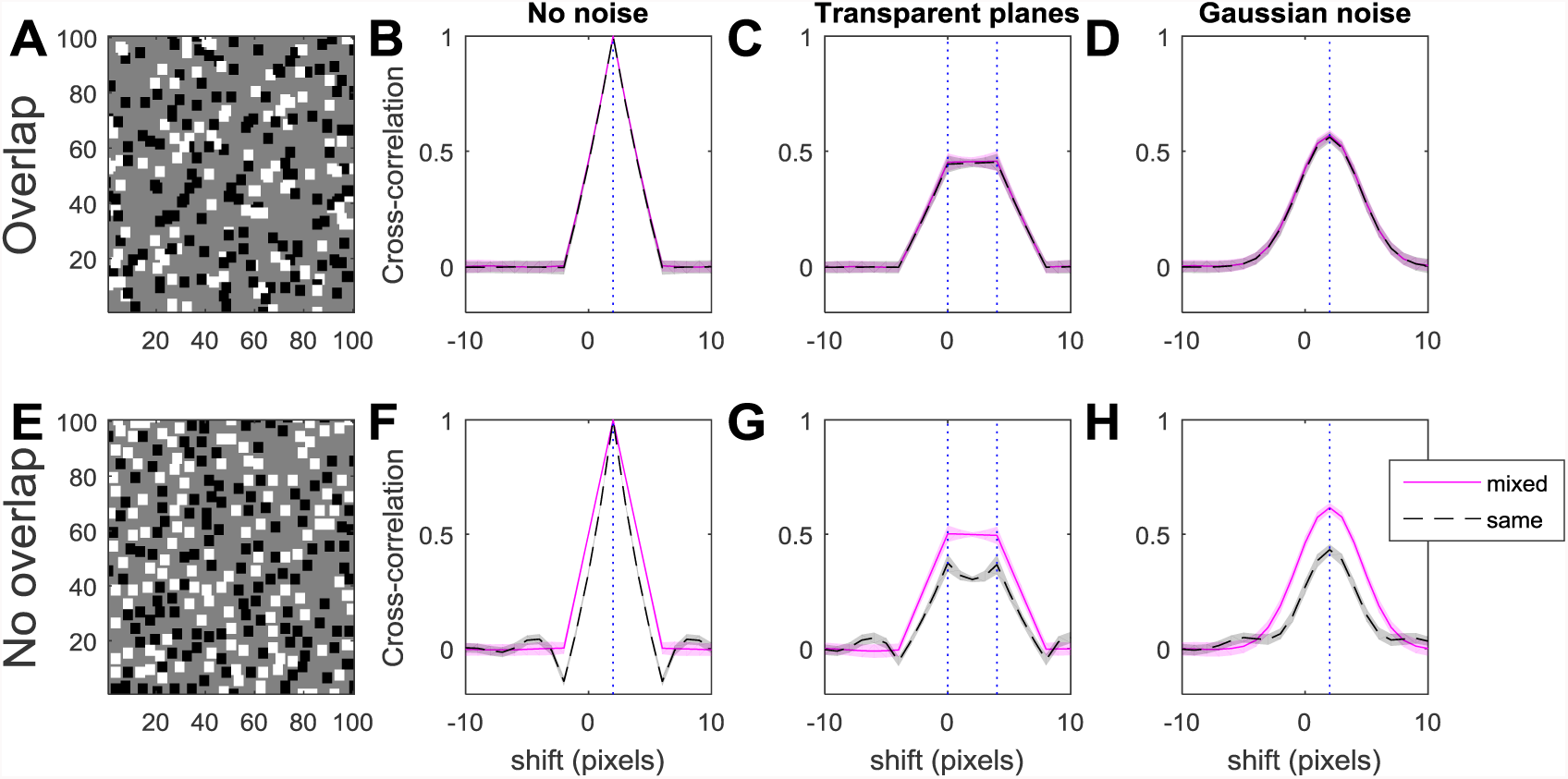
Cross-correlation functions between left and right images, for different types of disparity noise. Pink solid curves: results for mixed-polarity dot patterns like those shown on the left; black dashed curves: for same-polarity patterns, where the dots are either all black or all white. Top row: for random-dot patterns where dots are scattered at random; bottom row: where dots are not allowed to overlap. BF: “No noise”: all dots have the same disparity, 2 pixels. This thus has the same shape as the auto-correlation function of the monocular images, shifted to the stimulus disparity. CG: “Transparent planes”: half the dots (at random) have disparity 0 pixels and half have disparity 4 pixels. DH: “Gaussian noise”: every dot has random disparity noise drawn from a Gaussian of standard deviation 2 pixels. The images are 100×100 pixels and the dot size is 4 pixels. The curves show the mean cross-correlation function averaged over 100 different random images; shaded regions show ±SD.

### Image generation

In each run of a simulation, a mixed-polarity image-pair was generated first, with a background of 0 and white and black dots of ±1. The absolute value was then taken to produce a same-polarity stereogram with white dots on a gray background, and then the sign was inverted to produce a same-polarity stereogram with black dots on a gray background (cf Figure 1). Dots were square and the term “dot size” in this paper refers to the side of the square. To generate the pattern, first the number of dots was specified such that if none of the dots overlapped, a proportion *ρ* of the image would be occupied by dots. Dots were then added one after another. First, the luminance of the dot was chosen, either black or white with equal probability. *x* and *y* coordinates of each dot were chosen from a uniform random distribution across the image. The appropriate disparity was then applied to the *x* coordinate. In the Overlap condition, the dot was then simply drawn at the resulting location, overwriting any pixels belonging to existing dots. In the No-overlap condition, we checked to see if this dot would overwrite any pixels belonging to existing dots. If it did, the dot was abandoned and a new one was chosen. This process was repeated until the desired number of dots had been placed.

The monocular images were then normalised to have zero mean luminance and unit variance. This is a simple way of representing low-level adaptation in the visual system. As Read et al (2011) showed, if this step is not applied, energy model neurons have much lower-amplitude disparity tuning to same-polarity images, even in the absence of noise, but this artefact cannot explain the psychophysical results, both because early adaptation removes such changes in overall luminance, and because empirically the mixed-polarity advantage persists even if this manipulation is applied to the psychophysical stimuli. A very similar manipulation is implied in the definition of the Pearson correlation coefficient *r* (Equation 1), which is unchanged when a constant luminance offset is added to both images or when both images are scaled.

For clarity, image parameters are given in each figure legend. In most figures, the images were 100×100 pixels, the mean disparity was 0 pixels and the dot size was 4 pixels. For the neuronal simulations shown in Figure 5, we aimed to reproduce the stimulus of Figure 5 of Read et al (2011). In that paper, the dots were circles with an area of 8 square arcmin and, scattered without overlap, occupied 0.28 of the stimulus area; the step size was ~3 arcmin and the noise ~2 arcmin (precise values differed between observers, depending on what was needed to bring their performance to around 75% correct on average). In our simulation, the images were 241×241 pixels and the dot density was the same as in the experiments of Read et al (2011), i.e. ρ=0.28. We took 1 pixel to represent 0.5 arcmin and made the dot size 6 pixels, stimulus disparity 6 pixels and disparity noise 4 pixels.

**Figure 5.**
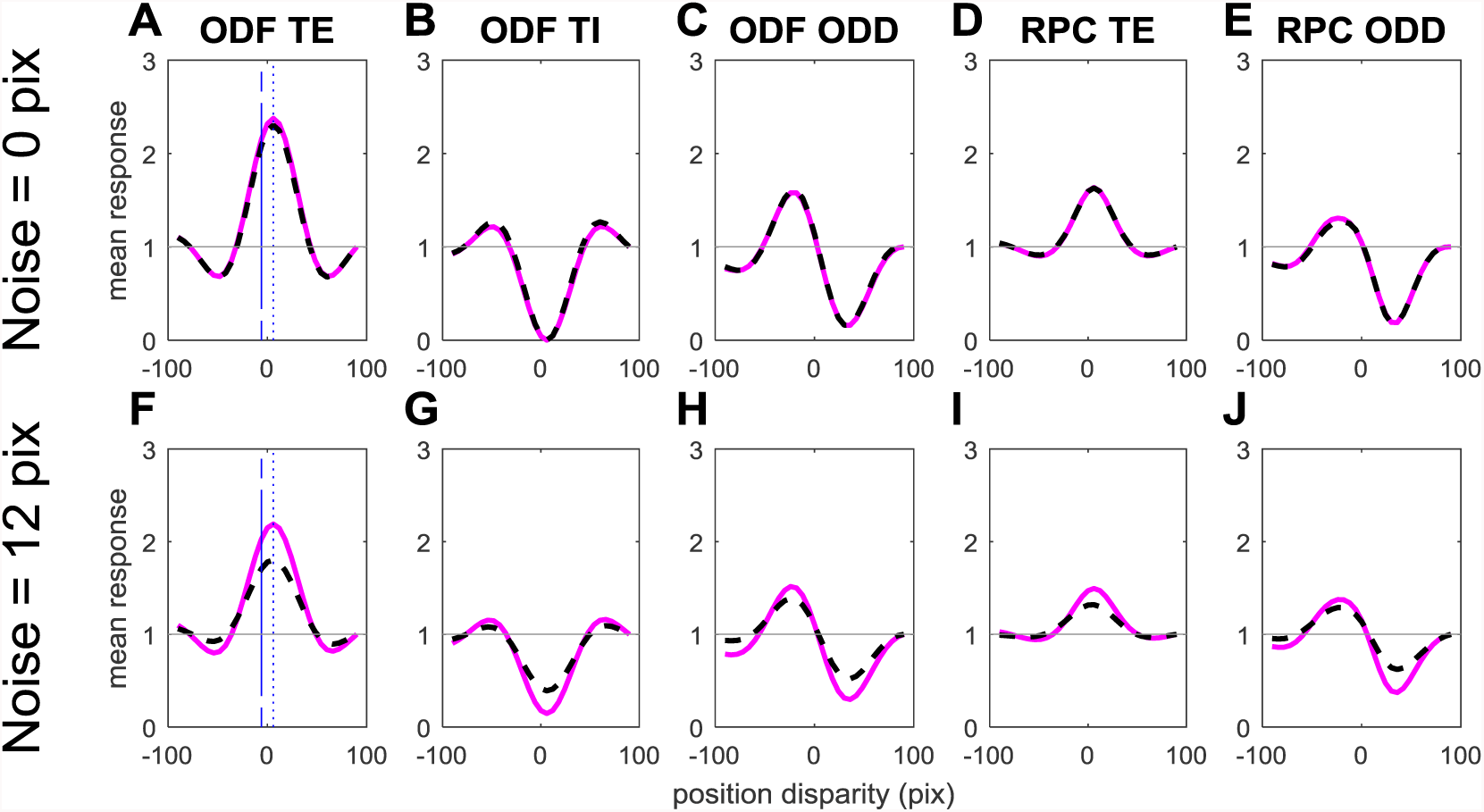
Population response for 5 different types of model V1 simple cells, for mixed-polarity (solid pink curve) and same-polarity (dashed, black) random-dot stereograms with no overlap. The stimulus mean disparity is 6 pixels. In the top row, the stimuli have no disparity noise, i.e. all dots have a disparity of 6 pixels. In the bottom row, each dot additionally has a noise disparity drawn from a Gaussian with a standard deviation of 12 pixels. The horizontal axis shows the position disparity of the neurons. Curves show the average response of a cell with the position disparity indicated on the horizontal axis to 10,000 random-dot stereograms with the specified parameters. The response of each neuron is normalised by its response to uncorrelated stereograms, marked with the horizontal gray line. The images were 241×241 pixels, dot size was 6 pixels, and images were normalized to have zero mean luminance and unit variance.

### Simulating response of binocular neurons

Our model receptive fields were even or odd Gabor functions:

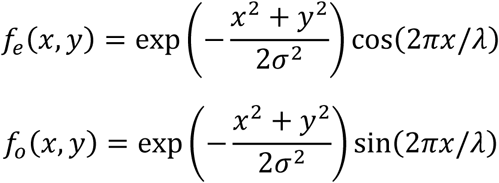

The receptive field standard deviation was σ=32 pixels, representing 16 arcmin or 0.27° since we are representing 1 arcmin with 2 pixels, and the carrier spatial period λ was 128 pixels or 1.1°. For a model neuron with position disparity *x*_0_ we computed the inner product of these each receptive fields, appropriately shifted, with the monocular images:

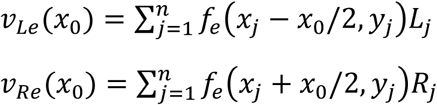

and similarly for v_Lo_. We considered various possible model V1 neurons, as follows:

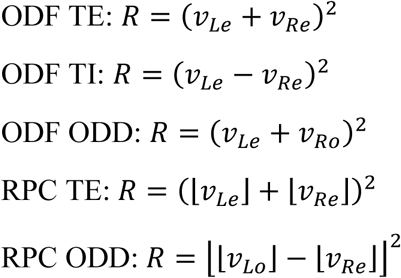

where the symbol └┘ denotes half-wave rectification, i.e. └*x*┘=*x* if *x*>0 and is 0 otherwise. Within each neuron class we simulated a population of neurons with different position disparities *x*_0_ (from −20 to +20 pixels in steps of 1 pixel). The tuning curves shown in Figure 5 represent each neuron’s mean response to 10,000 different random-dot patterns with the same disparity, normalised by that neuron’s mean response to binocularly uncorrelated stimuli.

“ODF” refers to the original form of the binocular energy model introduced by Ohzawa, DeAngelis and Freeman (Ohzawa, DeAngelis, & Freeman, 1990). “RPC” refers to the modified version introduced by Read, Parker & Cumming (2002), in which the monocular inputs are halfwave-rectified prior to binocular combination. “TE” denotes tuned-excitatory, i.e. a cell whose monocular receptive fields have the same phase and whose disparity tuning curve is therefore symmetric about a central peak (Read & Cumming, 2004). “TI” denotes tuned-inhibitory, i.e. a cell whose monocular receptive fields have opposite phase and whose disparity tuning curve is therefore symmetric about a central trough. “ODD” denotes a cell whose monocular receptive fields are π/2 out of phase. In the ODF model, such a cell has odd-symmetric tuning.

For simplicity, the example tuning curves in Figure 5 are all for simple cells. The same average tuning curves are obtained if we use phase-invariant complex cells, as for example *R* = (*v*_*Le*_ + *v*_*Re*_)^2^ + (*v*_*Lo*_ + *v*_*Ro*_)^2^, but complex cells show less variability in their response across different random-dot patterns.

## Results

### Where dots do not overlap, same-polarity stereograms have lower correlation than mixed-polarity

Figure 2 shows how the interocular image correlation declines as noise is added to the stimulus. The plots show the mean Pearson correlation coefficient between left and right images of random-dot stereograms, averaged over different random-dot patterns. The pink/solid lines show results for mixed-polarity stimuli, with equal numbers of black and white dots; the gray/dashed lines show results for same-polarity stimuli, where the dots are all black or all white. In the left-hand column, the patterns were generated such that dots are not allowed to overlap one another, as was done in Harris & Parker (1995); in the right-hand column, dots are scattered at random, with dots occluding others where they overlap. The top row is for a low dot density and the bottom row for a high dot density.

Unsurprisingly, image correlation falls as disparity noise increases. Where dots are scattered at random (Overlap condition, Figure 2BD), the decline is equal for mixed-polarity and same-polarity stimuli: the same amount of disparity noise results in the same decrease in correlation for both. However, where dots are placed so as to avoid overlap, the decline is steeper for same-polarity stimuli. This means that for a given amount of disparity noise, image correlation is lower for all-black or all-white dots. This effect becomes stronger as dot density increases (compare Figure 2C vs Figure 2A). Of course, where dot density is low enough, overlap will be rare anyway and so results must tend to be the same whether or not overlap is forbidden.

### The same effect applies for two shades of the same polarity

Harris & Parker (1995) also examined stimuli containing two dot colors but both with the same contrast polarity relative to the background, e.g. black and dark-gray dots on a gray background. They found that performance in this case was no better than for patterns containing dots of only one contrast. This result is also predicted by the image correlation. Figure 3 shows how image correlation is disrupted by disparity noise for the mixed- and same-polarity patterns considered before, and for “dark and darker” same-polarity patterns. Clearly the addition of “dark and darker” dots makes little difference; disparity noise is just as disruptive as for the other same-polarity patterns, and mixed-polarity patterns still have an advantage.

### Why forbidding dot overlap reduces image correlation

Why does this happen? Preventing dot overlap changes the pairwise statistics of pixels in the image. Consider a random-dot stereogram consisting of only black dots. Let *p* be the probability that a pixel in this image is covered by a dot and thus colored black; the probability that the pixel is grey background is then (1-*p*). If the dots are scattered randomly and independently, sometimes overlapping, then the probability *p* is uniform across the image. But this is not the case when overlap is forbidden. Consider a grey pixel adjacent to an existing dot, and consider all the possible positions of the next random dot that could turn this pixel black. Half of these random dot placements would overlap the existing dot, and hence are not permitted. As a result, the probability that this particular pixel is black is lower than the mean value across the image, *p*. This is reflected in the autocorrelation of monocular images. When overlap is allowed, the autocorrelation function is a triangle function (reflecting the autocorrelation function of a single dot), as shown in Figure 4B. But when overlap is not allowed, there are regions of negative correlation at displacements just larger than the dot width, caused by the reduced probability of black dots, as shown by the black dashed lines in Figure 4F. Note however that in mixed-polarity images the auto-correlation function is still a triangle function (pink curves in Figure 4F). This is because, while pixels adjacent to an existing dot are more likely to be grey, the probability of their being white or black is reduced equally, so that the mean correlation is unaffected.

The auto-correlation function, or the binocular cross-correlation where there is no noise, has a peak at 1.0 for all images (Figure 4BF). To understand what happens when noise is added, it is useful to consider a simplified case where the dots are given one of two disparities, depicting two transparent planes. Figure 4CG shows the cross-correlation for this situation; the blue dotted lines mark the disparities of the two planes. The images are the sum of two noise-free stereograms, for each of which the cross-correlation would be as shown in Figure 4BF, but shifted. Although the cross-correlation of the combined stereograms is not necessarily the mean of the cross-correlations for the individual stereograms, it closely resembles this. As illustrated in Figure 4G, for same-polarity stereograms the negative side-lobes in the auto-correlation reduce the peak cross-correlation, when there are multiple disparities.

Finally, Figure 4DH shows the cross-correlation function for stereograms with Gaussian disparity noise. The same effect applies, meaning that for No-Overlap random-dot patterns the cross-correlation function peaks at a lower value for same-polarity images than for mixed-polarity. This is the effect we saw in Figure 2C, where we plotted this peak correlation as a function of disparity noise.

### Standard models can explain the psychophysics

As we have seen, same-polarity stereograms end up with lower interocular correlation than mixed-polarity images affected by the same disparity noise. Human stereo performance declines as interocular correlation falls (Cormack, Stevenson, & Schor, 1991; Tyler & Julesz, 1978). It is therefore not surprising that humans perform better for noisy mixed-polarity stereograms than for same-polarity stereograms with the same amount of noise. Standard models of disparity encoding, such as the binocular energy model (Allenmark & Read, 2011; Filippini & Banks, 2009; Ohzawa et al., 1990; Qian & Zhu, 1997), are also based on interocular correlation, so we would expect these neurons to show stronger disparity tuning for noisy same-polarity than mixed-polarity stimuli.

Figure 5 confirms this expectation. It shows the mean response of a population of model V1 neurons to mixed-polarity (pink, solid curves) and same-polarity (black, dashed). In the top row, the stimuli are noise-free stereograms with a uniform disparity of 6 pixels, and the disparity tuning curves are identical for mixed- and same-polarity stimuli. This is the result reported by Read et al (2011) in their Fig 12. However, in the bottom row, we consider noisy stereograms, with the same mean disparity of 6 pixels but now disparity noise of 12 pixels. The noise effectively decorrelates the stimuli seen by the receptive fields, so the amplitude of disparity tuning falls for both stimulus types. However, as we have seen, the effective image correlation is lower for the same-polarity stimuli. Accordingly, the amplitude of disparity tuning for same-polarity stimuli is approximately half that for mixed-polarity.

Most obvious ways of decoding this population will therefore predict a mixed-polarity advantage in performance. For example, to simulate a front/back discrimination task, we could assume that the observer answers correctly when the response of the neuron tuned to the stimulus disparity of +6 pixels exceeds that of the “anti-neuron” tuned to −6 pixels (these neurons are marked with blue lines in Figure 5AF). The quantitative neurometric performance depends on the size of the receptive fields and the number of subunits. For our ODF TE simple cells, whose Gabor receptive fields have a standard deviation of 32 pixels or 5.3 times the dot size, we obtain a performance of 73% correct for mixed-polarity stimuli and 63% for same-polarity stimuli, giving an efficiency ratio of 3.6 (eq. 4 of Read et al 2011). For a pair of ODF TE complex cells, which sum input from pairs of subunits in quadrature phase, performance is better since the phase-independence reduces the image-dependent variability: 83% correct for mixed-polarity stimuli and 70% for same-polarity stimuli, an efficiency ratio of 3.3. These values are comparable to those for human observers. This confirms that the mixed-polarity advantage can be explained very well even by a single pair of classic energy-model simple cells. A complex neural network is not required.

## Discussion

For over twenty years, we have been puzzled by the evidence apparently supporting independent neural mechanisms for bright and dark information – ON and OFF channels – in human stereopsis. This evidence seemed to show that disparity was easier to extract in mixed-polarity random-dot stereograms – made up of black and white dots on a gray background – than in same-polarity stereograms, where all dots are black or all white. This conclusion is puzzling for two reasons. First, linear filtering in the early visual system means that even single-polarity random-dot stereograms stimulate both ON and OFF channels. Second, by the time disparity is encoded in the primary visual cortex, the evidence suggests that ON and OFF channels have been combined. Simulations confirmed that our current models predicted the same amplitude of disparity tuning for mixed-polarity as for same-polarity stereograms (Read et al 2011).

In this paper, we have shown that all previous results can be explained by a subtle stimulus artefact. It turns out that due to the way the stimuli were generated – preventing dot overlap – a given amount of disparity noise or of decorrelation, applied to individual dots, has a greater effect on the correlation of the entire image when dots all have the same contrast polarity relative to the background. We show that this effect produces a good quantitative account of human performance. This means that the mixed-polarity advantage is entirely consistent with our current understanding of disparity encoding and with other aspects of stereo psychophysics. Stereo correspondence is not, after all, intrinsically easier in mixed-polarity stereograms; it is simply easier in stereograms with higher interocular correlation.

If the stimuli are generated differently, with dots scattered at random and occluding one another where they overlap, then the image correlation ends up being the same for mixed- and for same-polarity stimuli. Significantly, the mixed-polarity advantage is then not observed (Read et al, 2011). This strongly suggests that the correlation artefact was indeed the reason for the mixed-polarity advantage in the original stimuli.

One of us (JCAR) has the dubious distinction of having examined the evidence for independent channels previously without having noticed this artefact. Read et al (2011) understood Harris & Parker’s result as implying that disparity is intrinsically easier to detect in mixed-polarity stimuli, and that noise functioned solely to remove the ceiling effect, bringing performance down to a level where this difference was observable. Accordingly, we examined the response of model neurons only to 100% correlated, noise-free stereograms, and concluded – wrongly – that these models could not account for the psychophysics. Goncalves & Welchman (2017) examined the response of their model to the actual, noisy stereograms used in experiments, and so revealed the effect which Read et al (2011) missed. Amusingly, Read et al (2011) even drew attention to the lack of dot overlap in Harris & Parker, and showed that the psychophysical mixed-polarity advantage was abolished if dots were allowed to overlap. However, we failed to appreciate the significance of this, attributing it to interference from the occlusion cue present in mixed-polarity stimuli.

We have now demonstrated that the psychophysical advantage for noisy mixed-polarity stereograms can be reproduced just as well with the standard binocular energy model as with the Binocular Neural Network of Goncalves & Welchman (2017). Both approaches judge depth based on which of two neurons, one tuned to near and one to far disparities, is responding most strongly. However, whereas Goncalves & Welchman’s decision neurons receive input from a convolutional network made up of thousands of binocular subunits, ours each receive input from just a single binocular subunit. This demonstrates that computation specific to Goncalves & Welchman’s Binocular Neural Network is not required for this result. We conclude that the differences in image correlation shown in Figure 2 are responsible for the mixed-polarity advantage in both models.

The mixed-polarity advantage was originally put forward as evidence that bright and dark information is processed separately in stereopsis, via distinct ON and OFF channels. We have now shown that the apparent advantage can be entirely explained by a correlation artefact in the stimulus. Of course, this result does not disprove the existence of independent ON and OFF channels in stereopsis, but it certainly undermines the current evidence for them.

